# Phase coding in phoneme processing slows with age

**DOI:** 10.1101/2023.12.22.573009

**Authors:** Konrad Dapper, Jakob Schirmer, Lukas Rüttiger, Etienne Gaudrain, Deniz Başkent, Sarah Verhulst, Christoph Braun, Ernst Dalhoff, Stehpan M. Wolpert, Marlies Knipper, Matthias H. J. Munk

**Author notes:** corresponding author’s.

## Abstract

The comprehension of phonemes is a fundamental component of speech processing which relies on both, temporal fine structure (TFS) and temporal envelope (TE) coding. EEG amplitude in response to phonemes has been identified as indicator of speech performance in hearing aid users. Presbyacusis may also alter neuro-electric responses to phonemes, even with minimally or unaffected hearing thresholds. Elevated speech reception thresholds (SRT) in absence of pure-tone threshold (PTT) elevation suggest central processing deficits. We therefore collected audiometric data (PTT, SRT) and EEG during passive listening in 80 subjects, ranging in age from 18 to 76 years. We confirm phoneme-evoked EEG response amplitude (PEA) as indicator of speech comprehension. Specifically, PEA decreased with elevated SRT, PTT and increased age. As novel observation, we report the temporal delay of phoneme-evoked EEG responses (PED) to increase with age and PTT. The absolute duration of PED, its age-correlation, and the lack of PEA-lateralization combined with the frequency of phoneme stimuli used here suggest a predominantly thalamic generator of phoneme-evoked EEG responses. Hearing loss in extended high-frequencies affects PED more than PEA. In our sample, neural compensation for increased PTT came at the cost of decreased temporal processing speed. Most importantly, PED correlates with SRT and explains SRT-variance in quiet and in ipsilateral noise that PTT cannot. PED was a better predictor of TFS coding in quiet and of TE coding in ipsilateral noise. As PED reflects both TFS and TE coding, thalamic activity may provide integrated information at the gate of neocortex.

**Significance Statement:** Intact speech comprehension is essential for social participation which protects against depression and dementia. Age-related hearing loss is a growing problem in aging societies, as hearing deficits constitute the third most important modifiable risk factor for cognitive decline. This work uses electrical brain responses to phonemes in a cohort covering age 18 to 76 years. As the temporal delay of phoneme responses showed the most significant correlations with age and high-frequency thresholds, we demonstrated that speed of neural processing seems essential for speech comprehension. The observed neural signals likely originate from thalamus which receives feedback from neocortex and is embedded in cognitive processing. Developing objective markers for speech processing is key for ensuring cognitive fitness in aging.

## INTRODUCTION

Age-related hearing loss (presbyacusis) is a growing problem in aging societies, particularly as it is associated with cognitive decline (Livingston et al., 2017; Gregory et al., 2020). The three main causes of presbyacusis are loss in cochlear-amplifier gain (Zelle et al., 2017), cochlear synaptopathy and loss of endocochlear potential (Keithley, 2020). Even though PTTs are the dominant factors determining SRTs (Wardenga et al., 2015) ,they can explain only approximately half of the total variance of SRT (Schirmer et al., companion paper (CoPa), also available on bioXriv). Age-related cochlear synaptopathy is a decrease in the number of auditory nerve fibers per inner hair cell during natural aging (Wu et al., 2019). Previous work suggests that cochlear synaptopathy and problems with speech intelligibility in the elderly may be directly related (Sergeyenko et al., 2013) and occur even without audiometric threshold elevation – at least within a frequency range of 6 kHz (Fullgrabe et al., 2014).

A prerequisite for the development of new therapeutic intervention is objective clinical testing of speech comprehension in humans with high sensitivity and specificity. For this goal we need to better understand the electrophysiology of speech encoding and its modification by age. Phonemes are the fundamental building blocks of speech. Understanding speech processing requires phoneme extraction from the continuous auditory stream, categorization due to prior experience and relaying information along the auditory pathway. Psychophysical experiments with vocoded speech, speech reduced to its temporal envelope (TE), demonstrated that speech content is predominantly encoded through amplitude modulation (TE coding) (Shannon et al., 1995). However, the brain can use temporal fine structure (TFS) for speech decoding even though TFS has been proposed to serve primarily the localization of competing sound sources (Smith et al., 2002), referring to the ability to switch between different speakers which has been shown to decrease with age (Meister et al., 2020). Identifying phonemes and words involves matching between sensory processing and language memory representations (Winkler et al., 1999; Altmann et al., 2014). Here, cortical and corticothalamic interactions are required (Hickok, 2012; Kegler et al., 2022) which is compatible with the findings that auditory brainstem responses (ABR) or auditory steady-state responses (ASSR) alone cannot fully explain speech comprehension beyond PTT (Guest et al., 2018; Carcagno and Plack, 2022; Schirmer et al., CoPa).

The summating-potential-to-action-potential ratio (SP/AP) is a measure for the number of auditory nerve fibers per inner hair cell. SP/AP is only weakly correlated with speech comprehension (Liberman et al., 2016; Grant et al., 2020; Chen et al., 2021) due to a large residual individual variance and many other contributing factors. One of the factors, which is certainly not represented by SP/AP, is cortical feedback (Hickok, 2012; Kegler et al., 2022) as speech comprehension does involve many cortical regions (Price, 2010; Moerel et al., 2014). Therefore, SP/AP is insufficient for clinical diagnoses of speech comprehension deficits. Consequently, a more general alternative based on advanced methods is required, which has yet to be developed.

One candidate for a more precise method to measure brain correlates of speech comprehension is EEG in response to individual phonemes and series of phonemes, revealing neural responses at middle and longer latencies (>6ms). It has been shown that the amplitude of speech EEG responses to syllables is a good predictor of speech comprehension in hearing aid users (Shetty and Puttabasappa, 2017).

Formants are a concentration of acoustic energy around a particular frequency in the speech spectrum (Titze et al., 2015). An important aspect of neural speech processing is addressing the simultaneously occurring redundant information present in the envelope of two or more distinct formants and the mechanism for combining formant information into unambiguous syllable representations, ultimately aiming at word identification through serial binding of subsequent syllables.

The present study demonstrates how delays in phoneme-evoked EEG responses can be quantified and utilized as a complementary observable to neuro-electric response amplitude, highlighting the importance of neuro-electric response delays for phoneme processing. We will discuss the functional localization of the phoneme EEG responses and the role of TFS and TE coding in the presence or absence of speech-shaped noise.

## Methods

### Participants

Eighty participants (18 to 76 years) were recruited after exhaustive information about the study. One participant had to be excluded because the earplugs for auditory stimulation fell off, and the participant did not inform the experimenters. The measurements analyzed for this report were part of a series of measurements comprising four sessions with audiometric, psychoacoustic, electro-, and magnetoencephalographic recordings. Participants were divided into three age groups, young (18-29 years), middle-aged (30-55 years) and elderly (56-76 years), which is not critical for the results of this report, but was used for illustration in Fig. 3. Details are given in the companion paper (Schirmer et al., CoPa). Before any assessment, audiological or physiological measurement, all participants gave written informed consent. The study design had been approved by the Tübingen University Hospital ethics board (392/2021BO2).

### Pure tone audiometry

The pure-tone threshold (PTT) measurements were performed by professional audiologists using the AT1000 Audiometer (Auritec, medizindiagnostische Geräte Gmbh, Hamburg, Germany). Pure tones were presented through Sennheiser HDA300 headphones (Sennheiser, Wedemark-Wennebostel, Germany) at the following frequencies (0.25, 0.5, 1, 1.5, 2, 4, 6, 8, and 10 kHz) and Sennheiser HDA300 at extended high frequencies of 11.2, 12.5, 14, and 16 kHz. Pure-tone averages over extended high frequencies (PTA-EHF; 11.3, 12.5, 14, and 16 kHz) and PTA4 (0.5, 1, 2, and 4 kHz) were derived from the right ear thresholds. These specific PTA groups were chosen to analyze thresholds for specific frequency ranges: (i) PTA4, because this is the worldwide standard for defining hearing-loss severity, (iii) (ii) PTA-EHF, because those frequencies display a possible role for speech perception (Hunter et al., 2020) and are above what is usually measured in a clinical setting.

### Speech audiometry

The well-established speech test in Germany is the Oldenburger Satztest (OLSA), a word-matrix based sentence test, in which the 50%-detection threshold is determined by adaptively adjusting the level of the presented sentences. (Wagener KC et al. Kollmeier B 1999, Zeitschrift für Audiologie/Acoustics; et al., 1999 ; Brand T and Kollmeier B The Journal of the Acoustical Society of America 2002), OLSA was performed in quiet and with speech-shaped noise at 70 dB SPL (ipsilateral) and with the same headphones (inserted earphones (ER1-14A) and mu-metal shielded ER2 Transducers (Etymotic Research, Elk Grove Village, Illinois, USA) as for the EEG experiment. For each noise condition, participants were presented with 20 sentences each, i.e. in unfiltered broadband noise (OLSA-BB), in low-pass filtered noise (OLSA-LP; frequency components >1500 Hz deleted from the OLSA power spectrum), or high-pass filtered speech (OLSA-HP, <1500 Hz; see Fig. 2 (Schirmer et al., CoPa)). Subjects were categorized according to their OLSA performance corrected for PTT influence by using a linear mixed model (Schirmer et al., CoPa).

### Phoneme stimuli for EEG recordings

Phoneme stimuli used for the EEG measurements were computer-generated using analysis-re-synthesis as implemented in the WORLD vocoder (Morise et al., 2016). They had a fundamental frequency of 116 Hz to match the OLSA sentences. The sounds /du/ (like in *Du*, “you” in German), /bu/ (like in *Budike*, “little shop”) and /o/ (like in *oder*, “or”) contained two formants below the phase-locking limit (PLL) in the human at ∼1500 Hz, while the phonemes /di/ (like in *die*, “the”), /bi/ (like in *bier*, “beer”) and /y/ (like in *üben*, “to practice”) contained only one formant below the PLL, which probably makes temporal fine-structure coding less important. TF-plotting of difference between phonemes pairs in Fig. 2 included a 1/f correction to compensate for the logarithmic frequency scale.

From each of the four stimulus pairs, a 9-step continuum was generated by gradually modifying the formants’ frequencies on a log-frequency scale. Their extreme contrasts were selected for the EEG experiment.

The stimuli were equalized such that the average level of the stimuli belonging to a given continuum was the same for all pairs, and was adjusted to 60 dB SPL (L_eq_). However, minor level fluctuations within a continuum were preserved to ensure that the level of the formants that remained identical throughout the continuum were not affected. Calibration was performed using a BK Type 4157 Microphone Brüel & Kjær (Böblinger Str. 13 · 71229 Leonberg, Germany ) in combination with an artificial ear with a volume of (1cm)³ and a 20 s integration time.

### EEG recording

EEG was recorded to study neural mid- and longer latency responses to phoneme stimuli (s. below). EEG recordings were accomplished with for passive stick-on electrodes (Neuroline 720, Ambu, Steinkopfstraße 4, Bad Nauheim, Germany) one mounted on each of the mastoids, the reference electrode on the center of the forehead below the participant’s hairline. The ground electrode was placed between the eyebrows. Signals were preamplified with a gain of 50 (EP50Amp) and digitized by actiCHamp Plus64 (Zeppelinstr.7 Gilching (Munich), Germany). The ADC rate was set to 50 k samples per channel.

All six phonemes (/du/, /bu/, /di/, /bi/, /o/, /y/) were presented in a frozen pseudo-random order. In total, we played 3456 presentations per stimulus of alternating polarity. In addition, we ensured that each transition between syllables was equally likely to minimize predictability. We presented 4 syllables per second jitterd in time by +/- 5 ms to minimize powerline noise via insert earphones (ER1-14A) and mu-metal shielded ER2 Transducers (Etymotic Research, Elk Grove Village, Illinois, USA). For the duration of the passive listening task, subjects laid on their back inside a soundproof chamber (Industrial Acoustics, Niederkruchten, Germany), while watching a silent movie (Metropolis) presented on a Tablet (Samsung a3) placed 60 cm above the subject’s head. The horizontal position was chosen to enable the subject to look straight ahead at the table without using their neck muscles. This way, subjects stayed awake during measurements. In case subjects fell asleep, they were immediately woken up by the experimenter. Those epochs were excluded from the analysis.

### Data analysis

Data analysis aimed at extracting amplitude and timing information from stable entrained neural responses to phonemes was achieved using MatLab (R2021b). First, all EEG data were resampled from the recording sample rate of 50k samples/second to 5 k samples/s with antialiasing at 4 kHz and a bandwidth of 200 Hz in order to speed up subsequent analyses. Then data were bandpass-filtered (60 - 2400 Hz) and segmented into 250 ms long epochs (-15 to 235 ms), including a 15 ms baseline. In order to minimize the influence of electrical stimulus artifacts at the level of EEG recordings, we averaged signals recorded with opposite polarity. Finally, all epoch-pairs with variance outside a region of three standard deviations around the mean epoch variance were rejected. Subsequently, the evoked signal’s continuous wavelet transform (cwt) was computed using the MatLab internal cwt function.

### Phase coherence

Three types of neural activity contribute simultaneously to the observable response: stimulus-evoked, stimulus-induced, and stimulus-independent signals. In the following, we will focus exclusively on stimulus-evoked responses. To maximize the signal-to-noise ratio in the investigated neural response, it is paramount to determine which time-frequency points are sufficiently dominated by the stimulus evoked response. For identifying the sufficiently evoked response components of the EEG signal, we determined phase coherence compiled for signals across different subjects by averaging the phase vector across all subjects. The averaged phase vector served as reference to which individual phase delays were compared. A time-frequency point was considered as coherent if the length of the observed phase coherence was more than 3.5 times larger than the chance level.

### Amplitude and delay of Phoneme evoked responses

Based on the phase-coherent points, two measures were extracted from the phoneme-evoked responses of each individual subject: phoneme-evoked delay (PED) and phoneme-evoked amplitude (PEA). PEA is the non-phase coherent average of all coherent time-frequency points computed as the average of the absolute values of the CWT coefficients. PED was determined as the average individual delay of all time-frequency points. Each individual time-frequency point delay is the phase difference between a subject’s evoked phase and the phase of the average recording across all subjects divided by the angular frequency.

### Pure tone normalized OLSA threshold (PNOT)

The primary factor that determines speech comprehension thresholds is the Pure-Tone threshold (PTT). However, other factors affect the OLSA threshold independent of PTT. To assess their effect on PED and PEA, the OLSA threshold needs to be corrected by PTT. In short, this normalization was achieved by performing least square multivariate regression (MatLab 2021b) between OLSA thresholds and the first five principle components (singular value decomposition, MatLab 2021b) of all PTTs. Next, the 5 PCAs of a single subject were used to evaluate the regression model resulting in a predicted OLSA value for each subject. For more details, refer to the companion paper (Schirmer et al., CoPa). Based on this analysis, we defined three groups of subjects stratifying them according to their PTT-normalized language comprehension performance: bad, standard, and good performers – for which we verified that all the groups had the same average pure-tone thresholds at all frequencies.

### Statistical evaluation

The differences between phoneme responses were assessed by z-transform with a standard deviation derived by 30,000 permutations of random phoneme pairs, ensuring that all subjects were present in both surrogate phoneme responses, thus ensuring that the reported difference is caused by the different stimuli and not by inter-individual differences. In order to evaluate correlations, a F-statistic against a constant model was used. The difference between the PNOT groups was achieved using a Kruskal Wallis test. Only after providing evidence for group differences by these non-parametric tests, regression was performed for parameters with significant group differences.

### Posthoc analysis of OLSA variance based on a linear mixed model

Analysis of variance for speech perception beyond pure-tone thresholds was performed in the same manner as in the companion paper (Schirmer et al., CoPa). The least-square multivariate linear fitting of the 5 PCAs derived from pure-tone thresholds and either PED or PEA was tested for its contribution to total speech-comprehension variance. To ensure uniqueness of the multivariate linear model, we first removed all linear correlations between the 5 PCAs and PEA and PED. This can be understood as removing the influence of PTT on PED and PEA. An inherent risk of the increase of dimensions of the regression model is the possibility of over-fitting. In order to eliminate this effect, we compared the observed increase in total explainable variance in the observed 6-dimensional model to the total explainable variance of 10.000 pseudo-models in which we randomly shuffled the additional observable before fitting the model. This gives us a reliable estimate of the gain in explained variance based on chance. The results are presented as stacked bar diagrams showing the percentage of variance that could be attributed to each of the observables. To better illustrate the magnitude of the effect, we computed the standard deviation, which can be attributed to the new observable. However, during this computation, we assumed a normal distribution of the 5 PCAs and the additional tested observable, an assumption that is not required for the statistical evaluation based on permutation analysis.

## Results

Our aim is to improve understanding of age-related hearing impairment, which is likely caused by cochlear synaptopathy in a significant proportion of elderly subjects. Cochlear synaptopathy, which refers to the loss of inner hair cell synapses with increasing age, has been hypothesized to be one potential cause of presbyacusis (Mohrle et al., 2016; Keithley, 2020; Tawfik et al., 2020). Based on findings by Shannon and colleagues showing that speech understanding is highly dependent on temporal envelope coding (Shannon et al., 1995), synaptopathy can be expected to alter syllable processing. In order to analyze neural correlates of phoneme processing, we performed speech EEG recordings in 80 subjects, which before underwent pure-tone audiometry (PTA4 and PTA-EHF) as well as speech audiometry in the form of the German matrix language test (OLSA). Our main results show that the neural delay of responses to phoneme stimuli provides a non-redundant measure of speech comprehension to the well-established pure-tone audiometric thresholds (PTT).

### Audiometric results

The most commonly used indicator of speech comprehension are pure-tone averages (PTA), which measures individual performance in two frequency ranges: PTA4 for speech-relevant frequencies between 0.5 and 4 kHz and the “extended high frequencies,” dubbed here as “PTA-EHF”, for frequencies between 11 and 16 kHz. Both thresholds, PTA4 and PTA-EHF, increase with age, and the slope of PTA-EHF thresholds across age was significantly steeper than the slope of PTA4 thresholds over age, indicating that EHF frequency regions of the cochlea were particularly sensitive for age-dependent hearing loss, as was previously described (Fletcher, 1950; Lee, 2013; Wu and Liberman, 2022). OLSA thresholds also increased with age. A more detailed analysis of the audiometric data of the same cohort is carefully documented in Schirmer et al. (Schirmer et al., CoPa).

### Phoneme-evoked EEG responses during passive listening

Combining audiometric tests like PTT and OLSA with speech EEG recordings is a powerful approach for unraveling subtle changes in neural processing induced by synaptopathy and presbyacusis. Speech-EEG was performed by recording the neuroelectric response during passive listening to six different syllables (/du/, /bu/, /di/, /bi/, /o/, /y/) while participants watched a silent movie in order to stabilize vigilance. All syllables evoked neural responses (**Fig.1**), which were analyzed by continuous wavelet transform (CWT) to extract stimulus phase-locked signal components. All responses cluster around the fundamental of the speaker’s voice (116 Hz) and to a lesser degree around its first harmonic (232 Hz) (Fig.1 A-F).

**Figure 1.**
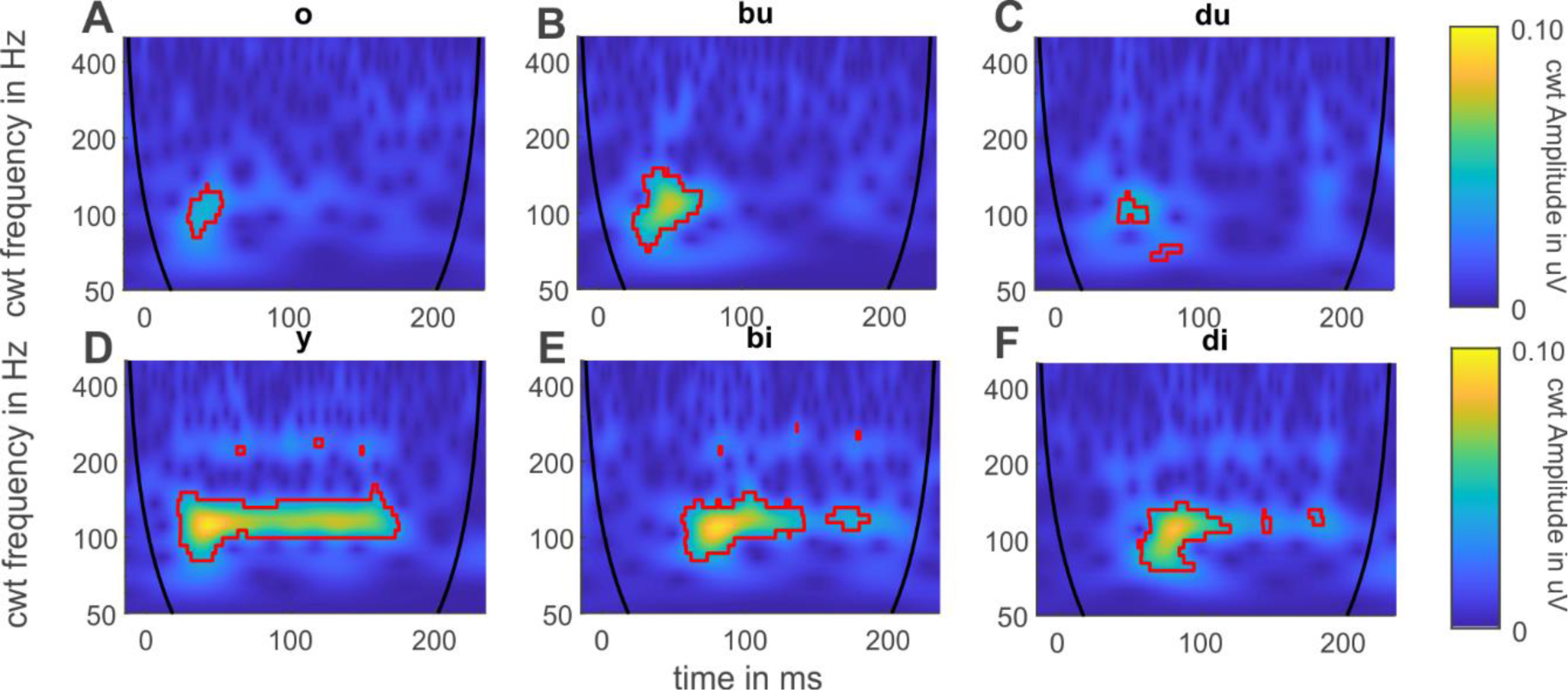
Averaged evoked EEG responses of all 80 participants (f/m/d 51/29/0) from both hemispheres during passive listening, presented in time-frequency space based on CWT. Black lines mark the cone of influence, indicating where edge effects (outside the lines in left and right lower corners) might occur in the CWT. Each plot A-F represents the response to a single phoneme, which was presented in pseudo-random order for a total of 3456 repetitions with two different electrical polarities. The x-axis represents time for each stimulus epoch in milliseconds; the y-axis shows frequency in Hz on a log scale; the z-axis represents amplitude in µV as pseudo-color; see color scales on the right. The red outline highlights all phase-coherent time-frequency points.

Syllables containing the vowels /i/ and /y/ (Fig.1 A-C) evoked stronger responses than syllables containing the vowels /o/ and /u/ (Fig.1 D-F). The phoneme /y/ evoked the most prominent responses (Fig.1A) while /o/ evoked the weakest (Fig. 1D).

### Phoneme evoked EEG responses differ depending on vowel-consonant contrasts and formants

As syllable discrimination is a prerequisite of speech comprehension (Guilleminot et al., 2023), we determined how the electric brain signals may carry information about differences of the tested syllables. Therefore, we compiled the z-transformed of pairwise amplitude differences of the respective EEG responses for combinations of syllables (**Fig.2, A-D**). These plots show that - at first glance - vowel contrasts cause larger brain electric differential responses than consonant contrasts as determined by a paired permutation analysis (compare bu-bi and du-di with bu-du and bi-di and see respective p-values in the legend). The lower plots (**Fig.2, E-H**) provide frequency-averaged z-scores over time, which were generated by integrating all z-values across all frequencies above 60 Hertz. These time courses illustrate that auditory stimulation (**Fig.2, I-L**) caused a consistent electric deflection, even if the response was weak as in the case of bi-di (**Fig.2 L**). Given the complexity of language decoding, we are not suggesting that simple linear combinations of stimulus frequencies predict response patterns and thus provide the brain with sufficient information for successful comprehension. As we will see later, interactions between peripherally distinct frequency channels seem to influence speech comprehension.

**Figure 2.**
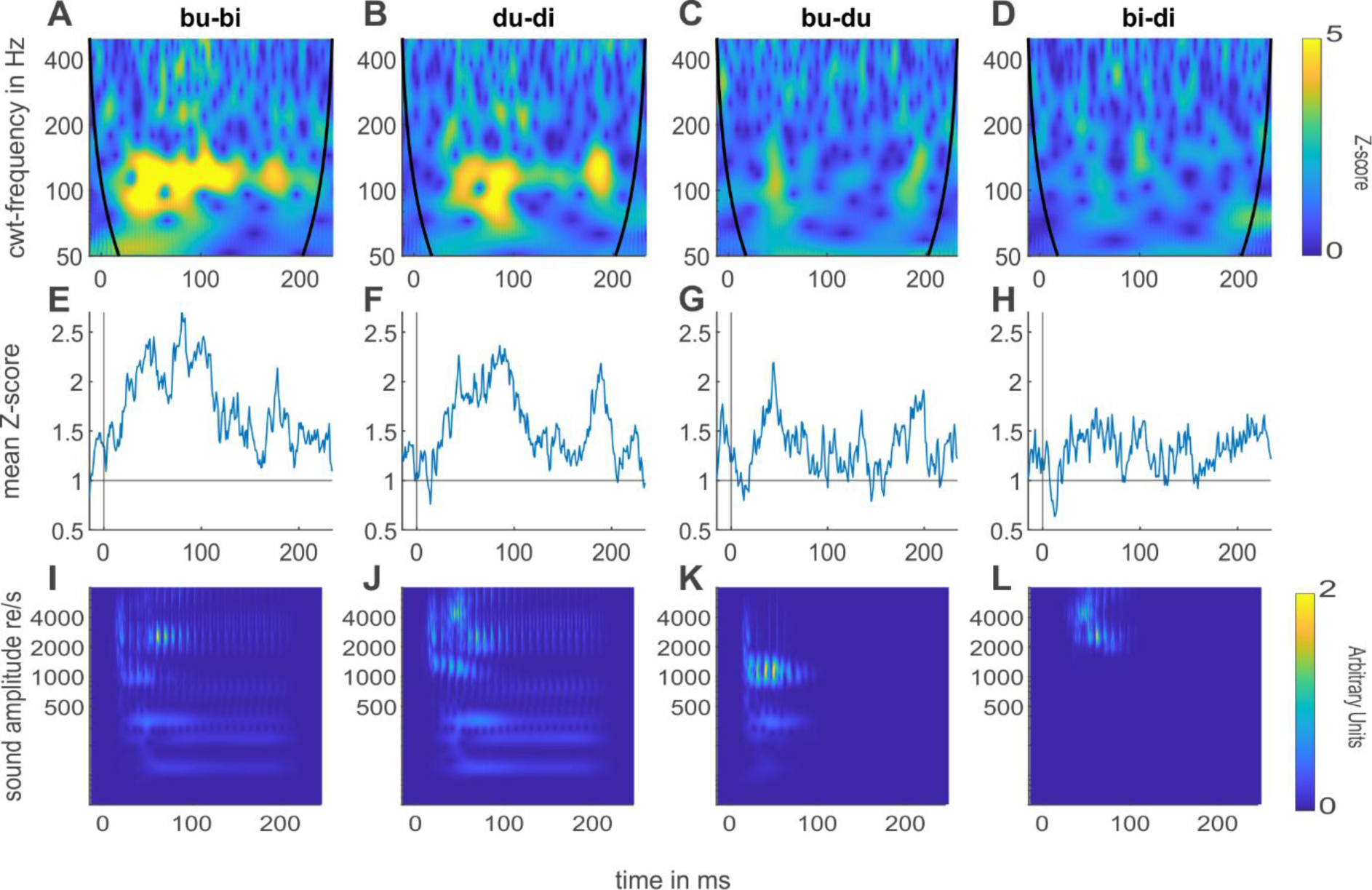
Electrophysiological difference responses for different pairs of syllables averaged across all subjects. **A-D**: permutation-based z-transform of response differences. **E-H**: time course of z-values averaged across all frequencies [60-2400 Hz]. Significance against random permutation of the time averaged z-score: p(bu-bi) < 1/30000 (***), p(du-di)<1/30000 (***), p(bu-du)=0.0174(*), p(bi-di)=0.1628 (n.s.).**I-L**: 1/f corrected TFA plots of phoneme stimuli differences. x-axis: time in ms; y-axis: frequency on log scale; z-axis: amplitude in arbitrary relative units.

### Phoneme-evoked EEG amplitude and delay correlate differentially with age, speech comprehension and hearing thresholds

In order to decompose EEG responses into complementary components, we next quantified amplitude (PEA) and delay (PED) and investigated whether these two very basic parameters would show any correlation with speech perception thresholds. In a first approach, we measured PEA and PED in individual recordings by averaging all time-frequency points which were at least 3.5 times above chance level for phase coherence (red outline in Fig.1). Both amplitude and delay are weakly correlated with the speech reception threshold (SRT) OLSA: low and high thresholds reflecting good and less good comprehension, respectively (**Fig. 3A, E**). Expectedly PEA decreased and PED increased with increasing speech comprehension thresholds. Similar effects could be observed for OLSA in noise (ipsi-and contralateral, not shown).

**Figure 3.**
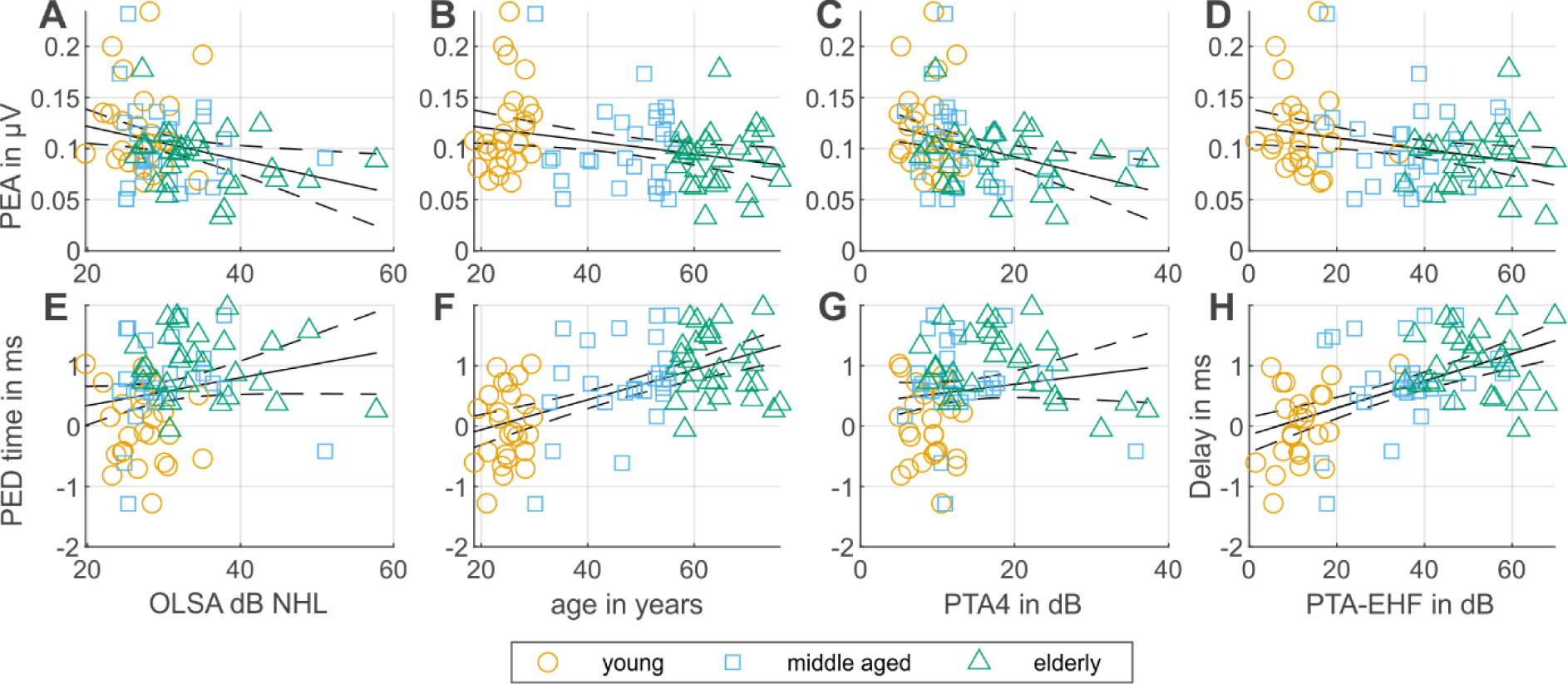
Age, OLSA and hearing thresholds affect PEA(A-D) and PED(E-H): **A-D** Speech-EEG amplitude and **E-H** delay as a function of age, PTA4, and PTA-EHF. Subjects are color-coded by their respective age groups: young (18-30y), middle-aged(30-56y), elderly (56-76y). Significance (F-statistic vs. constant model): p(amp/OLSA) = 0.01417(*),p(amp/age)=0.009(**), p(amp/PTA4)=0.004(**), (amp/PTA-EHF)=0.020(*), p(delay/age)=2.3e-8(****), p_delay_PTA4= 0.18(ns), p(delay/PTA-EHF)=5.8e-8(****), p(delay/OLSA) = 0.067823 (n.s.).

We then investigated correlations of PEA and PED with age (**Fig.3 B, F**) and pure-tone thresholds PTA-4 (**Fig.3 C, G**) and PTA-EHF (**Fig.3 D, H**). We observed several differences between delay and amplitude in these correlations: the two strongest and most significant correlations occurred for delay and age (p=2.3E10^-8^) as well as for PTA-EHF (p=5.8E10^-8^). The weakest correlation occurred with delay and PTA-4 (p=0.18 n.s.). In sharp contrast, the correlation between amplitude and PTA-4 (p=0.004) was more pronounced than the correlation between age and amplitude (p=0.009) and PTA-EHF and amplitude (p=0.02).

Taken together these observations show that the PED is largest in older participants with poor speech comprehension (Figure 3F, green triangles) that also exhibit highest PTA-EHF thresholds (Figure 3H, green triangles) while younger listeners have shorter PED (Figure 3F, yellow circles) and lower PTA (Figure 3G, yellow circles ) and exhibit lower PTA-EHF thresholds (Figure 3 H, yellow circles). PED and PEA are correlated but non-redundant as they explain only 18% of each other’s variance.

### Influence of consonants, vowels and formant frequencies on phoneme processing and how it is affected by speech comprehension performance

Our stimuli contained different combinations of vowels and consonants (s. Fig.2). Are the correlations observed in Fig. 3 driven by particular consonants or vowels, or, are those stimulus-independent features of auditory phoneme processing? Therefore, we plotted the regression lines for PEA and PED for responses evoked by syllables with the consonant /b/ and /d/ (Fig.4 A and E) and the regression lines for phonemes with two or only one formant below the PLL (Fig.4 B and F). Significant correlations of PEA with audiometric thresholds (OLSA, PTA4, PTA-EHF) were mostly negative, with one exception, the PEA of syllable pair /du-bu/ appeared unaffected by thresholds. Conversely, correlations with PED were almost all positive, but /di-du/ and /di-bi/ showed no positive correlation for PTA4 and only weak for correlations with OLSA. Typically, when PEA was negatively correlated with the audiometric thresholds, PED was either not or weakly positively correlated, while when PEA (s. e.g. /bu-du/) was not correlated, the correlation of PED with thresholds was particularly strong. This observation was more obvious in the correlation patterns with OLSA and PTA4, while correlations between PED and PTA-EHF appeared strong, independently of the correlations with PEA.

**Figure 4.**
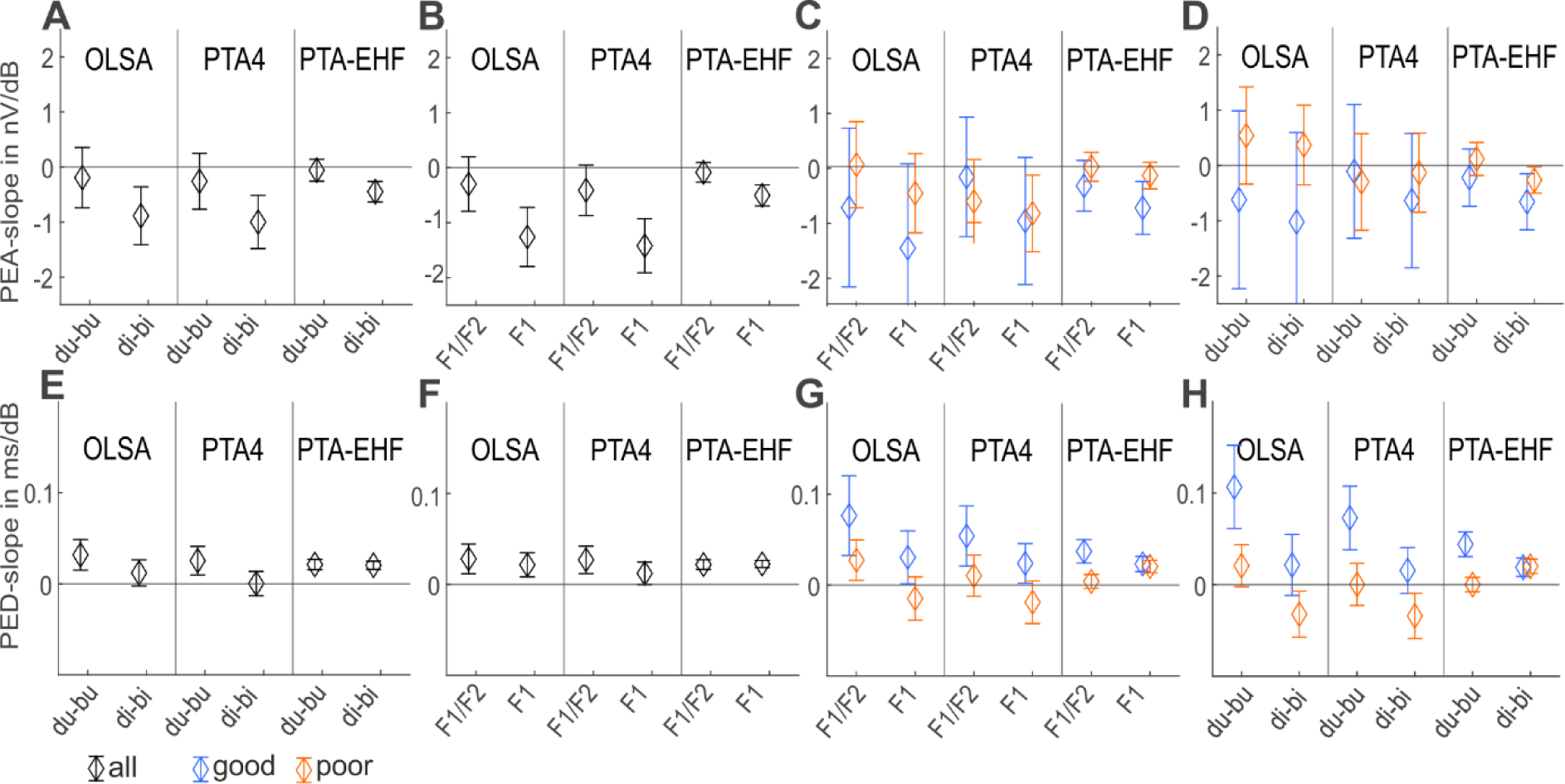
The slope of regression as a function of phoneme contrasts for thresholds in OLSA, PTA4, and PTA-EHF for one and two formants (B, F) and phoneme contrasts (A, E). **Upper row**: PEA [µV], **Lower row**: PED [ms]. Error bars label the distribution of fit errors of regression lines, for all subjects (n=80, black symbols), for a subsample with enhanced speech comprehension (n=26, blue symbols in C, D, G, H), and for a subsample with reduced speech comprehension (n=26, red symbols in C, D, G, H).

In order to differentiate neural processing mechanisms, we used phoneme stimuli with different formant properties: one group of stimuli (/di/ /bi/ /y/) contained only F1 as a formant below the phase-locking limit (PLL) (s. Tab. 1) and thus no redundant temporal fine structure (TFS) information was present, therefore TFS was limited for these phonemes. The other group of stimuli (/du/ /bu/ /o/) contained two formants F1 and F2 below the PLL (s. Tab. 1), i.e. temporal envelope information was redundant between the two formants. We will address the former as F1 and the latter as F1+F2 phonemes (s. Tab. 1). Thus, to improve the signal-to-noise ratio, we combined the responses of syllables with the same number of formants below the PLL when investigating the effect of redundant coding in the frequency range which in principle allows for TFS coding. Correlation analysis revealed how these neural phoneme responses relate to OLSA, PTA4, and PTA-EHF thresholds: Figure 4 B/F represents the slopes of the regression lines for PEA and PED with different audiometric thresholds, respectively. PEA was not correlated when two formants were present below the PLL (Fig.4B), while there was a negative correlation for phoneme responses with only one formant below the PLL (Fig.4B). In contrast, PED was – if at all - always positively correlated (Fig.4F).

**Tab1.** The table provides a comprehensive breakdown of the frequencies of the first four formants for each individual vowel sound, measured in Hz.

|  | /o/ | /du-bu/ | /di-bi/ | /y/ |
| --- | --- | --- | --- | --- |
| F1 | 380 | 200 | 350 | 350 |
| F2 | 750 | 750 | 1950 | 1780 |
| F3 | 3500 | 3500 | 2800 | 2100 |
| F4 | 10000 | 10000 | 4000 | 3000 |

As our main goal is to unravel mechanisms of phoneme processing relevant for speech comprehension, in particular speech comprehension beyond what can be explained with PTT, we defined groups based on PTA-normalized OLSA thresholds (dubbed “PNOT”, see methods). As can be seen in Fig.4C, PEA does not systematically co-vary with any of the thresholds, except perhaps PTA-EHF. In contrast, PED is almost always more strongly correlated in the good language comprehension performers (blue Fig.4 G/H), the notable exceptions being that the correlation between PTA-EHF and F2 phonemes, /di-du/ and /di-bi/ (see also Fig. 4 H), do not differ between good and bad comprehension performers.

### Progress towards an objective marker for speech comprehension deficits

Finally, in order to test whether individual PEA and PED values might be sufficiently informative about language understanding, we compared and ranked the correlations among PEA and PED with PTT-corrected SRTs with Kruskal Wallis tests and subsequent false-discovery rate correction. We found that the average PED of all syllables was significantly different among the three groups (good, indifferent and poor) of speech comprehension performers (p=0.004), while PEA was not (p=0.544), see **Fig. 5 A and C**.

**Figure 5.**
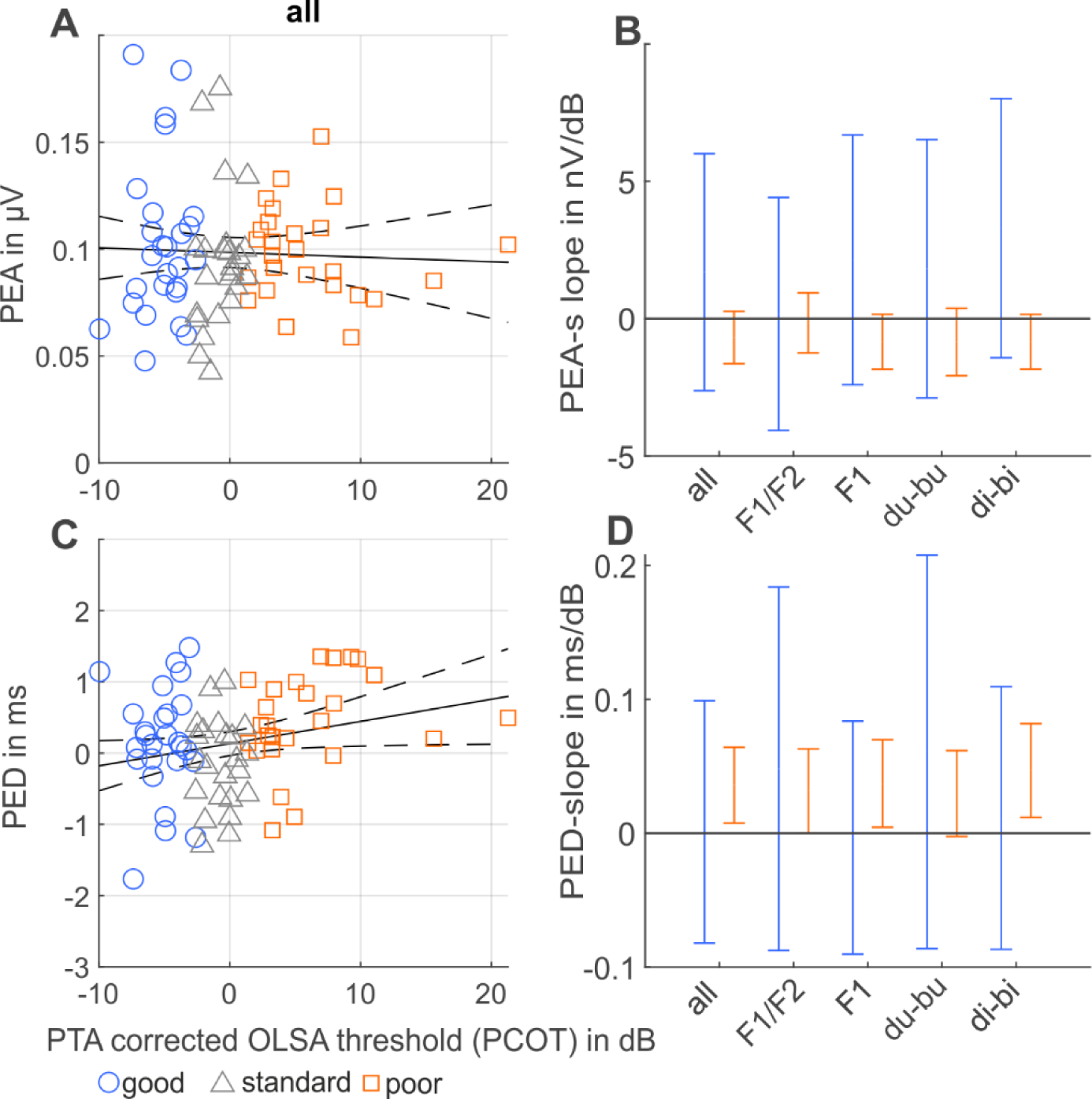
PTT-normalized speech comprehension thresholds (PNOT) do not affect amplitude (A), but do affect delay (C): **A** PEA and **C** PED as a function of PNOT for all phonemes. Subjects are color-coded with respect to their respective speech comprehension groups (blue=good, grey=standard, red=bad) based on PTA-corrected OLSA thresholds. **B, D** Slope of regression lines of PEA and PED with PNOT for speech perception groups, color convention as in A/C. Groups on x-axis: all phonemes, F1- and F2-phonemes, bi/bu, di/du, du/bu, di/bi, all split for good and bad performers.

The relation of PED and PNOT could for all syllables be confirmed via regression analysis with subsequent testing with an f-statistic (p=0.046) against a constant model. With the exception of one subgroup of phonemes (/di-bi/), all other phoneme groups returned significant PNOT-group (for bad and good performers) differences for the distribution of PED values, while for PEA, not a single significant result was obtained. Posthoc testing of individual group differences with rank sum tests revealed that there were significant differences between indifferent and bad performers, while good and indifferent performers showed only sporadic tendencies to be different.

Therefore, we next derived slopes (**Fig. 5 B and D**) and y-intersections (not shown) for all regression lines of the correlations between PEA and PED with PNOT for various subgroups of phoneme responses. For PEA (Fig. 5B) we found that the good performers showed no correlation, while the bad performers showed a systematic tendency towards negative correlations. For PED, we found the reverse: good performers again showed no correlation, while bad performers showed a systematic tendency towards positive correlations. In combination with the findings presented in Fig. 4, our observations suggest that listeners with good speech comprehension ability can either have a low OLSA threshold and fast processing as evidenced by short PED, or, have a high OLSA threshold combined with long PED and therefore slower phoneme processing, suggesting that the latter use the longer time interval for reaching successful phoneme discrimination. In contrast, subjects with poor speech comprehension performance who reached a low OLSA threshold had already long PED, while subjects with high OLSA threshold also had long PEDs which they could not prolong for compensation.

PNOT is derived from the three different OLSA frequency conditions (broadband, low- and high-pass), thus, it is not optimized for investigating different speech coding mechanisms as OLSA-BB provides acoustic information for both TFS and TE coding, while OLSA-LP only provides TFS information and OLSA-HP provides exclusively TE information. This could be rectified by looking at the amount of variance explained by PED for each of the 3 OLSA filter conditions independently. PED (Fig.6) does significantly contribute to explainable variance in speech comprehension, while PEA does not (data not shown). For OLSA in quiet, we found that PED explains variance of OLSA-BB (2% of variance or 1.5 dB in SRT) and OLSA-LP (3.6% of variance or 1.3 dB), but did not contribute to OLSA-HP (Fig.6A). This difference was more pronounced for PED caused by phonemes with two formants (Fig.6B) below the PLL (1.7 / 4.3% and 0.8 / 1.4 dB for BB/LP, respectively), while PED caused by phonemes with only one formant below the PLL (Fig.6C) showed only tendencies. In contrast, OLSA with ipsilateral noise, PED was predictive of OLSA-HP performance (5.8% 1.6 dB) and the difference was caused by phonemes with only one formant below the PLL (6.4%; 1.7 dB).

**Figure 6.**
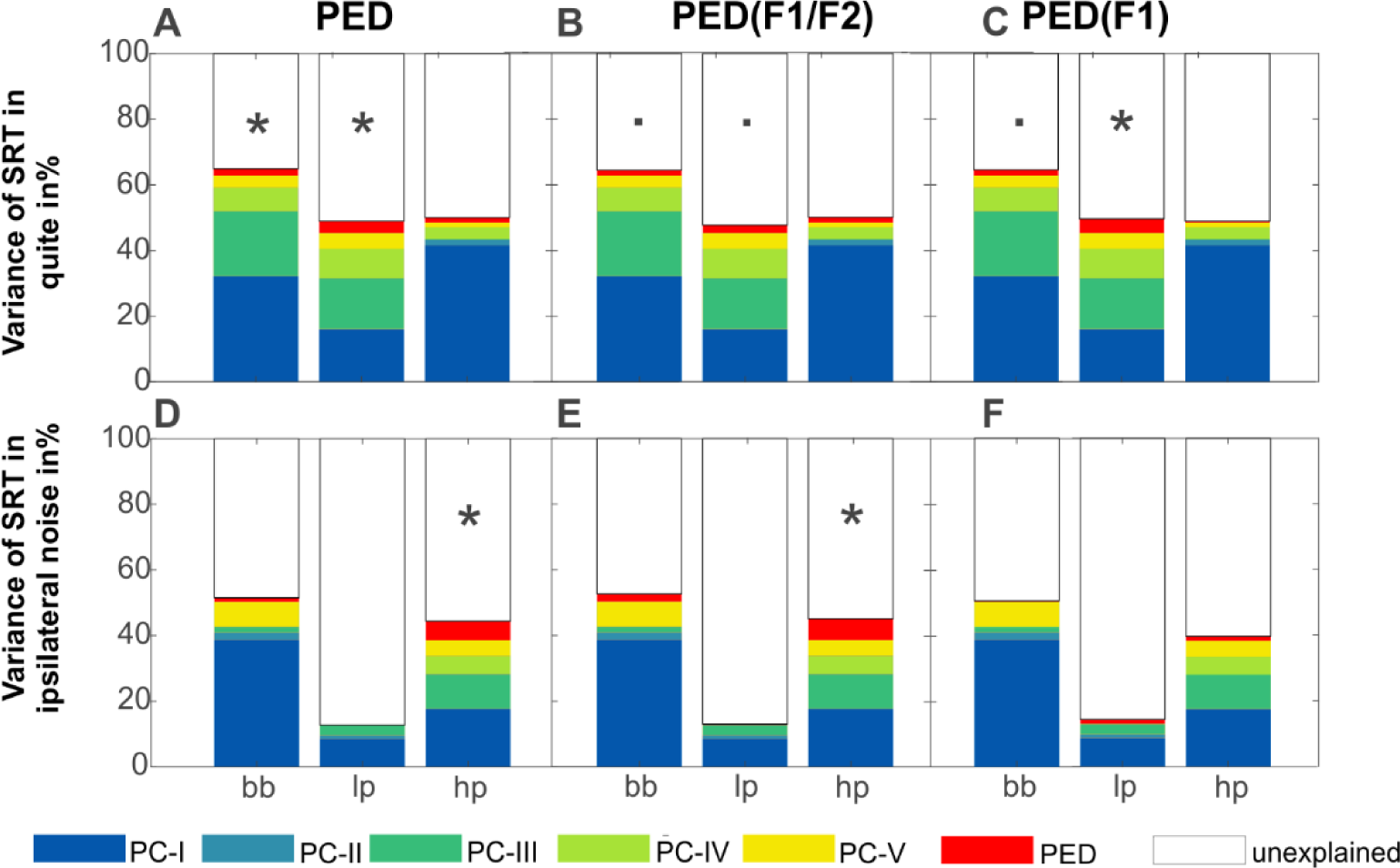
Variance explained by the first five PCAs of the PTT and PED for the 3 different OLSA conditions (bb=broadband, lp=low pass, hp=high pass). **A-C** results for OLSA in quiet, **D-F** results for OLSA in ipsilateral speech-shaped noise. PED was compiled for **(A, D)** the responses to all phonemes, **(B, E)** to phonemes with only one formant below the PLL (/di/, /bi/, /y/), and **(C, F)** to phonemes with two formants below the PLL (/du/, /bu/, /o/).

## Discussion

Neuro-electric responses to phoneme stimuli weaken and slow down with age and elevating pure-tone threshold (PTT). Our main observation is that the temporal delay of phoneme-evoked responses (PED) is an objective neural correlate of speech-comprehension deficits, even when speech-reception thresholds (SRTs) are corrected for PTT (Fig. 5 and 6). We thus extended prior findings on response amplitude in speech EEG (PEA) of hearing-aid users (Shetty and Puttabasappa, 2017) to listeners with no or only mild hearing impairment and in a wide age range. Our results highlight the importance of temporal delays in neuroelectric responses for speech comprehension.

On the one hand, in quiet, the z-transform of the difference between individual phoneme responses is larger when TFS differences were present in the phoneme stimuli. This is consistent with PED being more predictive for speech in quiet relying on TFS coding. On the other hand, when ipsilateral noise is present, PED is correlated to speech relying on TE coding, suggesting that PED integrates TFS and TE coding. Therefore, speech comprehension in quiet relies more on TFS coding, while in noise more on TE coding. This is further supported by the findings in our companion paper (Schirmer et al., CoPa) showing that more variance of SRTs is explained in the low-pass filtered speech-in-quiet condition with amplitude and latency of ABR wave I. Our quantitative neural findings on SRTs beyond PTT extend observations possible with ABR and ASSR responses, which have been shown to be at most weakly correlated with SRTs (Carcagno and Plack, 2022; Schirmer et al., CoPa).The effect was stronger when looking at delays rather than amplitudes ((Schirmer et al., CoPa), their figures 3 and 10B, C). One possible explanation for why speech-comprehension deficits are harder to observe at the level of neural brainstem response amplitudes is that ABR/ASSR and more than likely PEA are obscured by subcortical compensation mechanisms (Mohrle et al., 2016). However, amplification can never improve SNR needed for processing in higher brain regions or recover delay shifts as both of these would violate causality. The small magnitude of amplitude differences between PNOT-groups is surprising, because it is known that age-induced cochlear synaptopathy, measured via the SP/AP, influences speech comprehension (Grant et al., 2020; Chen et al., 2021).

Localizing the generators of speech EEG was not feasible at the level of dipoles due to having only two recording channels (Picton et al., 1999). However, we can infer an approximate location based on the physiological profile of the neural responses analyzed: a first factor, we want to consider here, is that ASSRs with high (>80 Hz) modulation frequencies reflect predominantly subcortical processes (Herdman et al., 2002; Lu et al., 2022). Therefore, the fundamental frequency of our phoneme stimuli (116 Hz) caused entrained EEG responses at frequencies associated with subcortical processing. A second factor to be considered for source localization is age-dependence of PEA as Ross and colleagues could show with source reconstruction of EEG and MEG recordings that the amplitude of cortical signal sources increases with age (Ross et al., 2020), while subcortical ASSR amplitudes decrease with age (Farahani et al., 2020). Our data show that PEA decreased with age (Fig.3B) again suggesting a predominantly subcortical origin of the signals described here. Furthermore, the absence of lateralization of neural phoneme responses recorded simultaneously at both mastoids also argues against a cortical source. Finally, to confine the potential source within subcortical regions, we consider the absolute magnitude of the delay prolongation with age which was about 2 ms.

This magnitude is consistent with the delay shift caused by different consonants in normal-hearing children (Johnson et al., 2008), but cannot be explained by the 400 µs age-induced ABR-wave-V shift in inferior colliculus (Konrad-Martin et al., 2012), or the longest ABR-latency-V difference between age groups in our cohort, that was more pronounced in participants with elevated PTA-EHF (Schirmer et al., CoPa).These observations suggest a supratentorial origin of the signal, which points to a thalamic origin of phoneme responses, likely coming from the MGN.

PED is more correlated with PTA-EHF than with PTA4 or SRTs (Fig. 3E-H). In contrast, PEA is more correlated with PTA4 and OLSA rather than PTA-EHF (Fig. 3A-D) demonstrating that delay and amplitude are related, but non-redundant observables. Beyond non-redundancy, PED allows for the investigation of causal processes in the ascending auditory pathway, which are instrumental for the time-critical segmentation of phonemes. Neural timing matters!

What does a prolonged PED mean for the mechanisms involved in language processing? We established that PED provides a new perspective on neural timing of speech coding. Some of the stimuli used here contain more than one formant below the human phase-locking limit (PLL) (Verschooten et al., 2019) providing redundant information in the frequency range supporting TFS coding. How does the presence or absence of redundant TFS information affect PEA and PED? On the one hand, when only one formant was present below the PLL, as in (/di/, /bi/, /y/), PEA consistently decreased with increasing SRTs, while PED was prolonged (Fig.5 and 6). On the other hand, PEA in response to phonemes with two formants below the PLL, as in (/du/, /bu/, /o/), is independent of SRTs as determined by OLSA, PTA4, and PTA-EHF, while PED was once again consistently prolonged with elevated thresholds, indicating that redundant information between formants could be used to recover PEA, but not PED. This is to be expected, because delays cannot be recovered without violating causality. PNOT did not systematically alter correlations between PEA and audiometric thresholds (Fig.5A, B). However, the group with impaired speech comprehension expressed systematically decreased correlation between PED and audiometric thresholds (Fig.5C, D). The largest effect occurred in the correlation of phoneme-evoked EEG responses to stimuli with two formants below PLL and the EHF-components. A dominant role of high-frequency signal components for speech comprehension could explain this.

In summary, our data show that PED was negatively correlated to OLSA, PTA and PTA-EHF thresholds. This finding is consistent with a model of speech comprehension that explains TE coding via saturation of high spontaneous firing rate auditory nerve fibers (high-SRANF) (Carney, 2018). High-SRANF are more prevalent in low-frequency regions of the cochlea (Bourien et al., 2014) their Fig.5B). Cochlear regions with more high-SRANF suffer proportionally more from fiber loss (Gleich et al., 2016), suggesting high-SRANF loss as a common driver between high frequency (e. g. PTA-EHF) threshold elevation and PED prolongation.

What is the role of TFS and TE coding for speech comprehension in the presence or absence of background noise and how is it reflected in neuroelectric delays? Our observed differences between neural phoneme contrasts are larger when TFS coding differences were present (as in /du/-/bu/) (**Fig. 2G, K**) compared to when they were absent (as in /di/-/bi/) (**Fig. 2H, L**). This relates to the view that speech comprehension predominantly requires TE coding (Shannon et al., 1995; Elhilali et al., 2009) and, to a lesser extent, TFS coding (Smith et al., 2002). The effect of background noise on different coding mechanisms based on the analysis of SRT variance shows that in quiet, PED is more correlated with OLSA-LP, which reflects TFS coding, while when measuring SRT in ipsilateral noise, PED is more correlated with OLSA-HP, which reflects TE coding. Therefore, PED likely measures neither exclusively TFS nor TE coding which may imply that the signal originating in thalamus already integrated both coding mechanisms.

Are neuroelectric phoneme responses influenced by cortical feedback? The neural difference response between bu and du (**Fig.2C**) persisted in time beyond the difference in their auditory input (**Fig.2K**). This suggests that the brain response, here depicted in the form of the integrated z-score (**Fig.2G**), reflects certainly more than mere feed-forward activity, most likely reflecting the expectancy of the listening subject, which is probably mediated by corticothalamic feedback (Brodski-Guerniero et al., 2017; Brodski-Guerniero et al., 2018) as these findings are not compatible with exclusive feed-forward-related TE coding. In light of our data, we have to revise the focus on the ascending auditory pathway when it comes to explaining speech comprehension deficits, because in our cohort a large part of SRT variance could not be explained by any of the peripheral auditory parameters, which is consistent with previous findings (Carcagno and Plack, 2022).

In conclusion, neural timing in the auditory system is - beyond response amplitude - a non-redundant feature for information coding, which slows with age in a way that contributes to speech comprehension beyond the effects of threshold elevation. As we have shown for the first time, presbyacusis not only affects PTA-EHF, but also prolongs the delay of entrained neural responses (PED) by up to 2 ms, in particular, when a second formant is present below the PLL. Together with the general envelope frequency preferences of subcortical and cortical circuits and lacking lateralization of the neural responses, our findings point to the thalamus as the most likely signal source. Furthermore, PED explained an admittedly small but significant amount of SRT-variance in quiet as well as in the presence of ipsilateral noise, suggesting that PED bears potential for new diagnostic procedures with which features of presbyacusis independent of PTT elevation might become objectively quantifiable.

## Acknowledgments

All measurements were performed as a Port of the ERA CoSySpeech project (BMBF 01EW2102 CoSySpeech)

